# Enabling *in vivo* Analysis Via Nanoparticle-mediated Intracellular Assay Probe Delivery: Using RAS as the Prototype

**DOI:** 10.1101/2020.07.05.188862

**Authors:** Fengqian Chen, Qi Liu, Terrell Hilliard, Tingzeng Wang, Ziye Dong, Wei Li, Hongjun Liang, Weimin Gao, Leaf Huang, Degeng Wang

## Abstract

Many experimental protocols must be executed *in vitro* due to a lack of cell-permeable analysis probes. For instance, the cellular signaling moderator RAS proteins alternate between the active GTP-binding and the inactive GDP-binding states. Though many GTP analogs can serve as probes for RAS activity analysis, their cell impermeability renders *in vivo* analysis impossible. On the other hand, the lipid/calcium/phosphate (LCP) nanoparticle has enabled efficient intracellular delivery of a nucleotide analog as a chemotherapy agent. Thus, using RAS analysis and LCP nanoparticle as the prototype, we tackled the cell-impermeability issue via nanoparticle-mediated intracellular delivery of the analysis probe. Briefly, BODIPY-FT-GTP-γ-S, a GTP analog that becomes fluorescent only upon protein binding, was chosen as the analysis probe, so that GTP binding can be quantified by fluorescent activity. BODIPY-FT-GTP-γ-S-loaded LCP-nanoparticle was synthesized for efficient intracellular BODIPY-FT-GTP-γ-S delivery. Binding of the delivered BODIPY-FT-GTP-γ-S to the RAS proteins were consistent with previously reported observations; the RAS GTP binding activity was reduced in serum-starved cells; and a transient activation peak of the binding activity was observed upon subsequent serum reactivation of the cells. In a word, nanoparticle-mediated probe delivery enabled an *in vivo* RAS analysis method. The approach should be applicable to a wide variety of analysis protocols.

## 1. Introduction

Many biochemical and molecular and cellular biology protocols rely on usage of analysis probes with quantifiable readout, such as colorization, fluorescence and specific antibody binding. Often, the probe is an analog of a key component of the targeted biochemical process, such as a reactant/product or an allosteric regulator of the enzyme, enabling direct assessment of how active the process is proceeding. For instance, the protein kinase assay uses ATP molecules with tagged γ-phosphate group to monitor the transfer of the group onto substrate proteins, *i*.*e*., the kinase-catalyzed phosphorylation reaction. As another example, the Global Run-on Sequencing (GRO-seq) analysis uses the bromo-tagged UTP (BrUTP) and its antibody to quantify genome-wide nascent RNA production (1).

Unfortunately, many such protocols have to be performed *in vitro* due to poor cell permeability of the analysis probes. For instance, both the ATP analogs with tagged γ-phosphate group and BrUTP discussed above are not sufficiently cell permeable, preventing *in vivo* kinase substrate tagging and efficient BrUTP incorporation into nascent RNA, respectively. One obvious drawback is the uncertainty about whether the *in vitro* results faithfully reflect the underlying process in living cells. Thus, *in vivo* protocols have always been vehemently pursued. For instance, the development of cell permeable and calcium sensitive fluorescent dye for *in vivo* calcium concentration measurement (2,3); the development of fluorescent proteins, such as green fluorescent protein (GFP), for protein expression and tracking in live cells (4,5); the use of fluorescence resonance energy transfer (FRET) in *in vivo* detection of protein-protein interaction (6,7); and the use of cell permeable 4-Thiouridine (4sU) in place of BrUTP in metabolic labeling of RNA molecules to quantify their production and stability (8-10). However, many important protocols remain exclusively *in vitro*, waiting for solutions to the membrane impermeability issue to enable the *in vivo* versions.

The study of the RAS small GTPases, for example, currently lacks *in vivo* methods (11). The RAS genes (K-RAS, H-RAS and N-RAS) were discovered and named for their central roles in sarcoma-virus-induced <u>ra</u>t <u>s</u>arcoma formation. They are well established as the most frequently mutated oncogenes in human cancers, and RAS-driven cancers are among the most difficult to treat (11-13). The underpinning of this clinical significance is RAS acting as a crucial hub of the cellular signaling network. RAS proteins receive signals from many upstream receptor tyrosine kinases and other signaling pathways. Their downstream effectors include PI3-kinase, RAF, RalGDS and many other key signaling molecules. By alternating between active GTP-binding and inactive GDP-binding states (14,15) (Fig. 1A), they coordinate upstream signals and downstream regulatory effectors to maintain optimized cellular harmony with the extracellular environment. Many GTP analogs are suitable to act as as probes for analyses of RAS and other G-proteins, but their cell impermeability make it impossible to do the analysis *in vivo* in live cells (16-21). Currently, RAS activity measurement methods are mostly *in vitro* (22). One popular approach is the GST-RBD pull-down assay – quantification of the amount of the RAS protein in the cell lysate that binds to a recombinant Glutathione S-transferases (GST) tagged RAS-binding domain (RBD) of a RAS effector protein, usually the RAFs.

**Figure 1.**
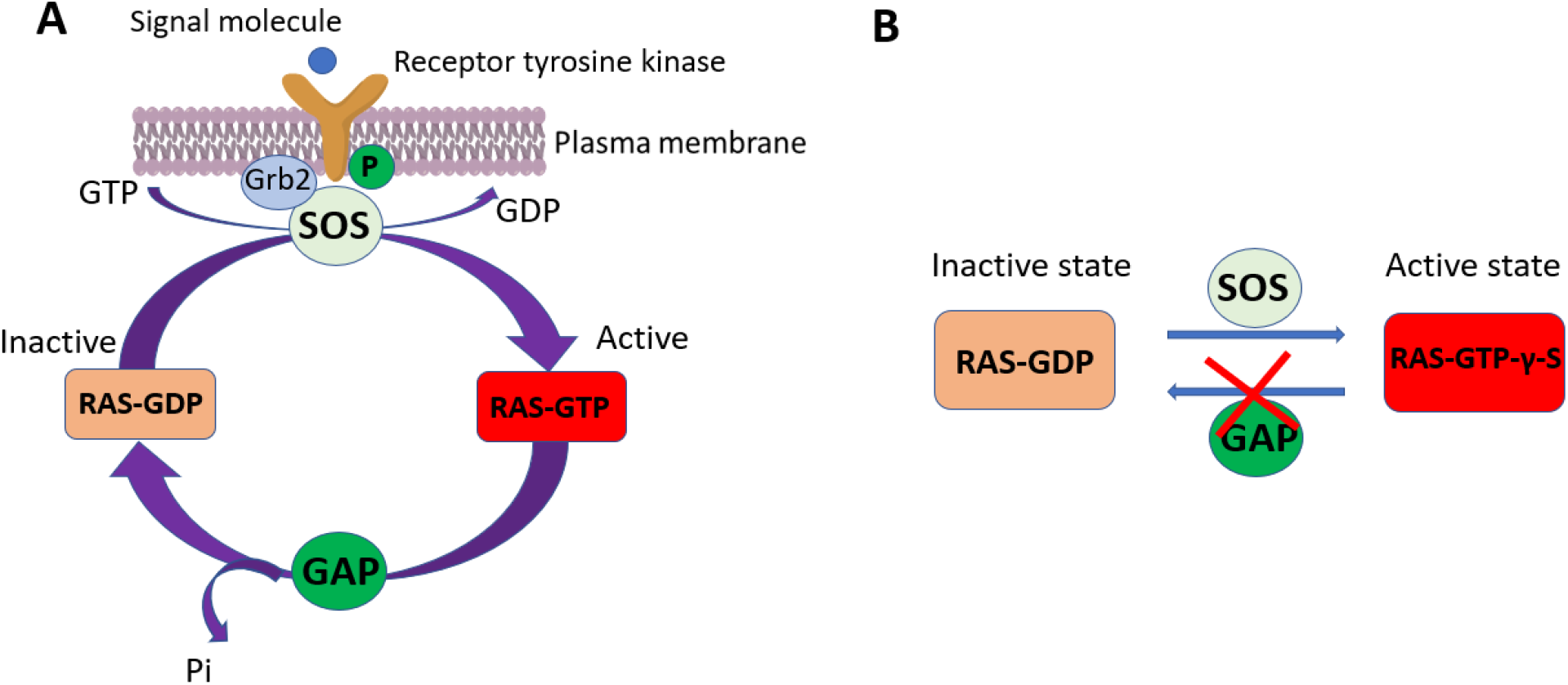
Schematic illustration of the RAS signaling cycle (A) and intervention by GTP-γ-S and its derivatives (B). (A) Upstream signals activate SOS, replacing the RAS-associated GDP with a GTP and activating RAS signaling to downstream effectors. The RAS GTPase activity then hydrolyzed the GTP into a GDP, shutting off downstream signaling. (B) Binding by the GTP analog GTP-γ-S, which is much less hydrolysable, leads to stabilization of the RAS GTP-bound active state. Thus, GTP-γ-S derivatives with quantifiable reporter activities, such as fluorescence, have been used as probes in *in vitro* analyses of RAS and G-proteins.

Nanoparticle drug delivery systems have, on the other hand, achieved great successes in overcoming this cell impermeability issue (23,24). It is temping to ask whether the nanoparticles are capable of delivering the *in vitro* analysis probes and, thus, enable *in vivo* versions of the analyses. Luckily, in case of RAS analysis, the lipid/calcium/phosphate (LCP) nanoparticle, one of the widely used drug delivery vehicles (25), provides an ideal opportunity to answer this question. It consists of a calcium phosphate (CaP) core, which carries the to-be-delivered chemicals, and a cell-specific lipid bilayer, which encapsulates the CaP core and enables cellular uptake of the nanoparticle through endocytosis. It has been used for efficient intracellular delivery of the chemotherapy drug Gemcitabine Triphosphate (26,27), a nucleotide analog (28). Since Gemcitabine Triphosphate is structurally similar to the GTP analogs used as probes in *in vitro* RAS analyses, it is plausible that the LCP nanoparticle is capable of efficient intracellular delivery of the GTP analogs. Thus, the LCP nanoparticle and the GTP analogs, together with the RAS analyses, form a testing ground for the principle that nanoparticle-mediated probe delivery can enable *in vivo* versions of previously *in vitro* analysis methods.

The analysis of RAS and other G-proteins often use GTPγS, a much less hydrolysable GTP analog due to the replacement of an O by an S in the γ-phosphate group, as the probe. As mentioned earlier, once bound to GTP, the RAS proteins are in the active state (Fig. 1A). When the GTP is hydrolyzed to GDP, they revert to the inactive state. Thus, GTPγS stabilizes RAS in active state, as shown in Figure 1B. To facilitate various biochemical analysis, GTPγS is often conjugated to a tag with quantifiable reporter activities such as fluorescence (18,20). One such GTPγS derivative is BODIPY-FL-GTP-γ-S, whose fluorescent activity is ∼90% quenched in solution but activated upon binding to a protein (19,21). This conditional fluorescent activity makes it a perfect probe for quantification of RAS GTP binding and, thus, signaling activity. However, as with many other nucleotide analogs, BODIPY-FL-GTP-γ-S cannot penetrate the cell membrane and get inside the cell independently.

Thus, in this study, BODIPY-FL-GTPγS-loaded LCP nanoparticles were synthesized for delivery into the human HCT116 cells. The delivery was efficient enough for the delivered BODIPY-FL-GTP-γ-S to compete with endogenous GTP for binding to the RAS proteins. The RAS binding activity was consistent with expectation based on previous observations. Thus, an alternative RAS analysis method based on *in vivo* RAS-GTP binding is established. Such repurposing of nanoparticle for intracellular delivery of analysis probes should be applicable to many more research protocols.

## 2. Materials and methods

### 2.1. Materials

Alex-Fluor-647-ATP, BODIPY-FL-GTPγS and the pan RAS monoclonal antibody (Ras10) were purchase from Thermo Scientific (New York, US). Dioleoyl phosphatydic acid (DOPA), 1,2-dimyristoyl-3-trimethylammonium-propane chloride salt (DOTAP), and 1,2-distearoyl-sn-glycero-3-phosphoethanolamine-N-[amino(polyethylene glycol)-2000] ammonium salt (DSPE-PEG_2000_) were obtained from Avanti Polar Lipids, Inc. (Alabama, US). DSPE-PEG-anisamide (AEAA) was synthesized in Dr. Leaf Huang’s lab as described previously (29). Other chemicals were purchased from Sigma-Aldrich, Inc. (Missouri, US).

### 2.2. Cell culture

HCT116 human colon cancer cells, originally obtained from Dr. Brattain’s lab (30-33), were cultured in the McCoy’s 5A medium (Invitrogen, USA) supplemented with 10% fetal bovine serum, 100 U/mL penicillin, and 100 mg/mL streptomycin (Invitrogen, USA). Cells were cultivated at 37 °C and 5 % CO_2_ in an incubator.

### 2.3. Preparation of LCP nanoparticles

The nanoparticle was synthesized as previously described (26,27,29). Briefly, 20 μL of 5 mM specified nucleotide analog solution (BODIPY-FL-GTPγS or Alexa-Fluor-647-ATP) in 12.5 mM Na_2_HPO_4_ solution and 100 μL of 2.5 M CaCl_2_ solution were dispersed in cyclohexane/Igepal solution (70/30, v/v), respectively, to form an oil phase. After 15 min of stirring, 100 μL of 25 mM DOPA was added into the oil phase for 15 min, then 40 mL of 100 % ethanol was added into the oil phase. Later, the mixture solution was centrifuged at 10,000 g for 20 min to pellet the LCP particle core out of the supernatant solution. After washing with 50 mL ethanol twice, 100 μL of 10 mg/mL cholesterol, 100 μL of 25 mg/mL DOTAP, 100 μL of 25 mg/ml DSPE-PEG, and 10 μL of 25 mg/mL DSPE-PEG-AEAA were added into the LCP core solution. After evaporating under N_2_, the residual lipids were suspended in PBS to produce the layer of LCP nanoparticle. After being sonicated for 10 min, the LCP solution was dialyzed in PBS.

### 2.4. Characterization of LCP nanoparticles

The sizes of LCP nanoparticles were determined with the Malvern dynamic light scattering (DLS) equipment (Royston, UK). The concentration of BODIPY-FL-GTPγS was determined by an HPLC spectrophotometer (Shimadzu Corp., Japan). Transmission electron microscope (TEM) photos of BODIPY-FL-GTPγS-loaded LCP nanoparticles were observed under JEOL 100CX II TEM (Tokyo, Japan).

### 2.5. Intracellular uptake analysis by flow cytometry and confocal fluorescent microscopy

Intracellular uptake of nanoparticles was monitored with synthesized Alex-Fluor-647-ATP-loaded LCP nanoparticle, using the Alex-Fluor-647 fluorescence as the probe. For flow cytometry analysis, log-phase HCT116 cells were treated with either the nanoparticle or PBS as a negative control. At specified time points, the cells were trypsinized and suspended into fresh medium, followed by three centrifugation-wash-suspension cycles. The suspended cells were then analyzed with an Attune NxT flow cytometer (Thermo, US) to quantify their intracellular fluorescence intensity. For microscope analysis, HCT116 cells were plated into glass-bottom confocal dishes (MatTek, US) and incubated for 24 h. Then, the cells were treated with the nanoparticle for 2.5 h with a concentration of 1 μg/mL. And the cell nucleuses were labeled with Hoechst 33342 dye for 15 min. Finally, the images were taken by a Zeiss 880 confocal microscope (Germany).

### 2.6. RAS immune-precipitation and fluorescence spectroscopy analyses

The cells were washed by cold PBS twice and then lysed with non-SDS lysis buffer that contains protease and phosphatase inhibitors for 30 minutes at 4 °C. Total protein concentrations of the cell lysates were determined with a BCA protein assay kit (Thermo Fisher Scientific, USA). The cell lysates were pre-cleaned by an off-target control antibody and aliquoted. One aliquot of the cleaned cell lysates were incubated with 10 μL anti-RAS monoclonal antibody under rotary agitation at 4 °C for 12 h. 100 μL SureBeads protein G magnetic beads (Bio-Rad, USA) were added. The cell lysate, antibody and bead mixture was incubated with rotary agitation at 4 °C for 4 h. The beads were precipitated and washed twice with wash buffer.

The beads were then transferred into a 96-well plate. The plate was analyzed with an Infinite M1000 Quadruple Monochromator Microplate Reader (Tecan) to quantify the fluorescent activity co-precipitated with the RAS proteins.

### 2.7. Western blot analysis

Another aliquot of the pre-cleaned cell lysate described above for immune-precipitation was used for Western blot analysis. The lysates were separated by SDS–PAGE, and then transferred onto a nitrocellulose membrane. After blocking the NC membrane with 5 % non-fat milk for 1 h at room temperature, it was incubated with the santi-RAS antibody at 1:1000 dilution at 4 °C overnight. Membranes were washed five times with TBST solution, incubated with the corresponding secondary antibody (1:3000 dilution) for 1.5 h at normal room temperature, and washed five times with TBST solution. The RAS protein bands were visualized using an enhanced chemiluminescence (ECL) system according to the manufacturer’s instructions (Thermo Fisher Scientific, USA).

## 3. Results

### 3.1. Synthesis and Characterization of LCP nanoparticles

As mentioned earlier, the LCP nanoparticle has been used for efficient intracellular delivery of the nucleotide analog Gemcitabine Triphosphate as a chemotherapy agent. Herein, we tested it for intracellular BODIPY-FL-GTPγS delivery to overcome the cell impermeability issue. Figure 2 shows a schematic illustration of the preparation procedure of BODIPY-FL-GTPγS LCP nanoparticles. BODIPY-FL-GTPγS is first loaded into the CaP core of the LCP nanoparticles, which is then coated with a DOPA layer. DOPA functions to prevent the aggregation of the CaP core because of the strong interaction between the CaP core and the phosphate head of DOPA. The cationic 1,2-dioleoyl-3-trimethylammonium-propane (DOTAP) lipid is then added as an outer layer onto the DOPA layer (27). Additionally, an outside layer of neutral dioleoylphosphatidylcholine (DOPC) with DSPE-PEG with a tethered targeting ligand anisamide (AA) is added. These layers keep the nanoparticles stable in the hydrophilic tissue culture medium and also enhance cellular uptake of the nanoparticles. Upon nanoparticle cellular internalization, the CaP core is rapidly dissolved in the endosome due to its low pH, leading to increased endosome osmotic pressure. Consequently, the endosome swells and ruptures, releasing the loaded BODIPY-FL-GTPγS into the cytoplasm. This quick process also minimizes lysosomal degradation after endocytosis.

**Figure 2.**
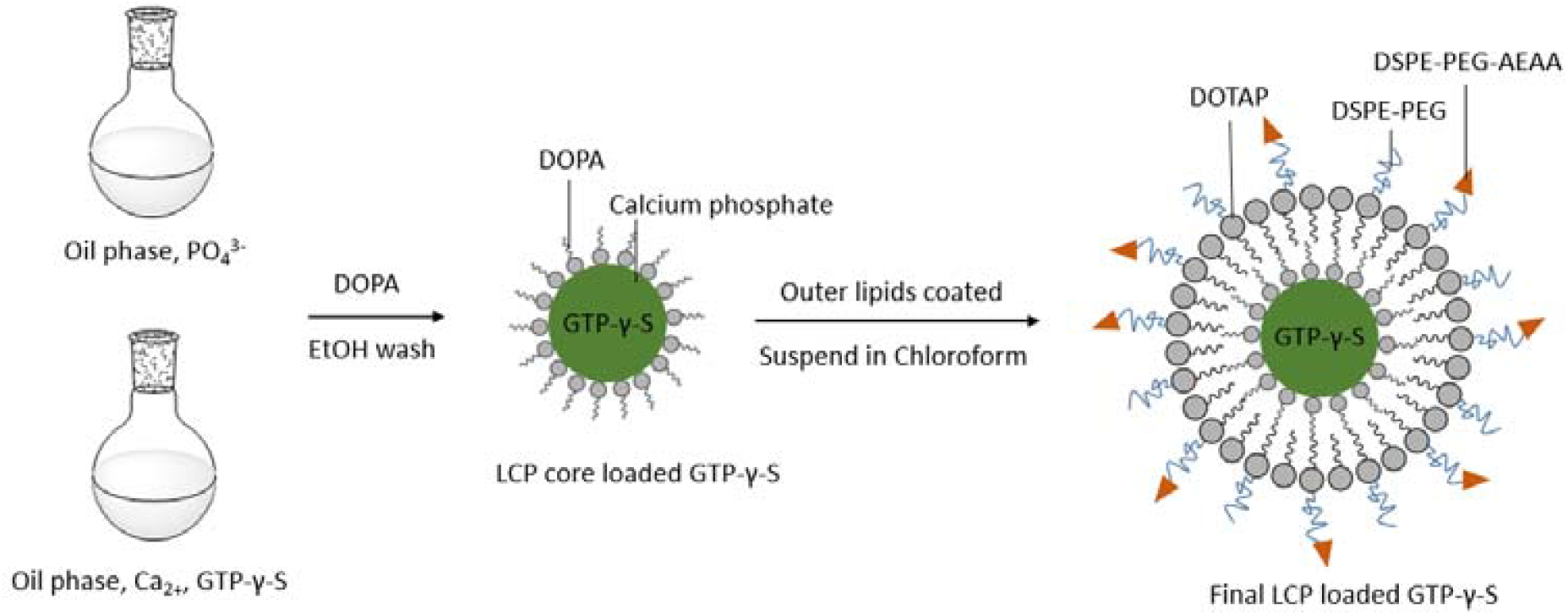
Schematic illustration of the synthesis procedure of BODIPY-FL-GTPγS-loaded LCP nanoparticle. Please see Materials and Methods for details.

Figure 3 summarized the results of our characterization of synthesized nanoparticles. The BODIPY-FL-GTPγS-loaded LCP nanoparticle sizes are exhibited in Figure 3A in a histogram. The sizes are also summarized in Figure 3C. On average, BODIPY-FL-GTPγS-loaded LCP nanoparticles were 124.6±7.1 nm. A transmission electron microscope (TEM) image is also shown in Figure 3B, illustrating the monodispersed nanoparticles with spherical shapes. The final concentration of BODIPY-FL-GTPγS in 5 mL of produced nanoparticles was 10 μM. Thus, about 50% of the starting BODIPY-FL-GTPγS (20 μL of 5 mM solution) was loaded into the nanoparticles.

**Figure 3.**
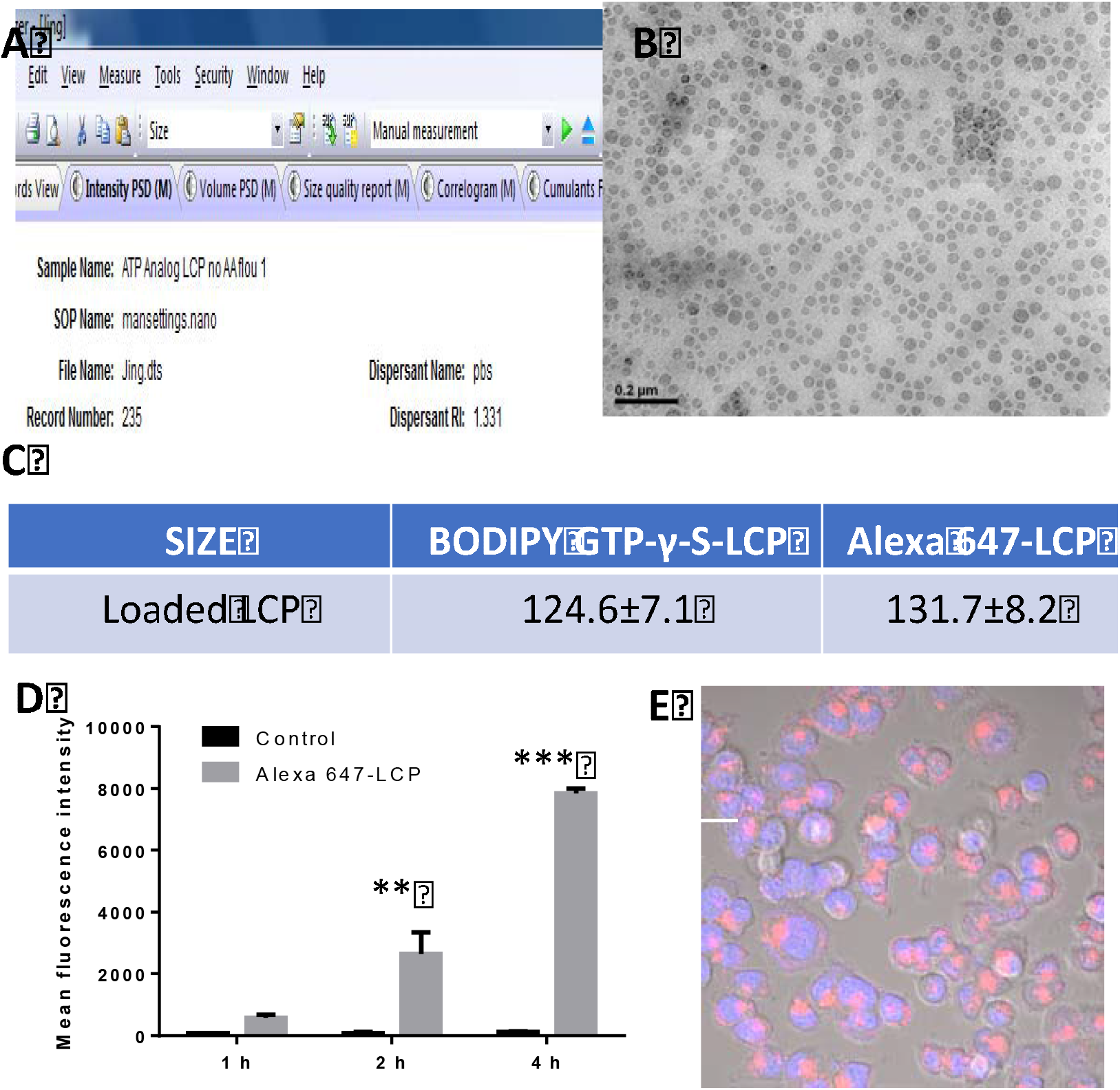
Characterization of the LCP nanoparticles. (A) Histogram of BODIPY-FL-GTPγS-loaded LCP nanoparticle size. (B) TEM images of BODIPY-FL-GTPγS-loaded LCP nanoparticles. The experiment was repeated 4 times. A representative result is shown. The scale bar is 200 nm. (C) Average sizes of BODIPY-FL-GTPγS-loaded LCP nanoparticles and Alexa-Fluor-647-ATP-loaded LCP nanoparticles. (D) Cellular uptake of Alexa-Fluor-647-ATP-loaded LCP nanoparticles in HCT116 cells monitored by flow-cytometry analysis. And (E) Fluorescent microscope image of HCT116 cells treated with Alexa-Fluor-647-ATP-loaded LCP nanoparticles. This experiment was repeated 4 times. A representative result is shown. The scale bar is 20.0 μm..

### 3.2 Cell uptake of LCP NPs

We next monitored intracellular uptake of the LCP nanoparticle. For this purpose, we used the fluorescent ATP analog Alexa-Fluor-647-ATP, so that the uptake can be conveniently tracked by fluorescent activity. This ATP analog was loaded, as described in Materials and Methods, into the LCP nanoparticles. On average, Alexa-Fluor-647-ATP-loaded LCP nanoparticles have a size of 131.7±8.2 nm. The synthesized nanoparticle was added into the tissue culture medium and incubated with the HCT116 cells for intracellular uptake. At 1, 2 and 4 hours, as described in Materials and Methods, the cells were harvested and analyzed with flow cytometer to measure their fluorescent activity. The result is shown in Figure 3D. Compared with control cells, nanoparticle treated cells exhibited significantly stronger fluorescent activity by 1 hour of the treatment, suggesting cellular uptake of the nanoparticles. While no appreciable increase of the fluorescent activity was observed in the control cells, the fluorescent activity kept increasing in the nanoparticle treated cells.

This uptake was also directly observed as described in Materials and Methods. The cells were grown in a glass bottom dishes and treated with the nanoparticle for 2.5 hours. They were then fixed, nucleus-stained and observed under a confocal microscope. As shown in figure 3E, strong Alexa-Fluor-647 red fluorescence activity was spotted inside the HCT116 cells, demonstrating the cellular internalization of loaded LCP nanoparticles. Thus, we were convinced that the LCP nanoparticles could be used as a vehicle for intracellular nucleotide analog delivery.

### 3.3 *In vivo* RAS binding to nanoparticle delivered BODIPY-FL-GTPγS

Encouraged by the cellular uptake of the nanoparticle, we explored the potential of using it to deliver fluorogenic probes to enable an *in vivo* version of a previously *in vitro* analysis protocol. An excellent candidate is the fluorogenic GTP analog BODIPY-FL-GTPγS. This molecule is not highly fluorescent in solution, but becomes activated upon binding to proteins such as RAS. We tested whether this fluorogenic property can be used to visualize general GTP-protein binding in live cells. Thus, we synthesized nanoparticle loaded with a mixture of BODIPY-FL-GTPγS and Alexa-fluo-647-ATP (10 to 1 molar ratio). The constitutively fluorescent alexa-fluo-647-ATP was used to monitor nanoparticle delivery; BODIPY-FL-GTPγS, to the contrary, becomes fluorescent only upon binding to proteins, and thus was used to monitor protein-GTP binding. Briefly, the fluorescent signals of the two molecules didn’t co-localize (Fig. 4B). The constitutive ATP analog is heavily concentrated inside the nanoparticle before its release and, thus, gave strong localized signal inside internalized nanoparticles (Fig. 4B, red signal). BODIPY-FL-GTPγS, on the other hand, was not fluorescent inside the nanoparticle and in its free form upon release from the nanoparticle. Instead, its fluorescent activity largely localized to the cell membrane (Fig. 4B, green signal), where G-proteins are known to be largely anchored to and thus GTP-protein binding events concentrate. As our control, cells treated with empty control nanoparticle did not exhibit the fluorescent signals (Fig. 4 D).

**Figure 4.**
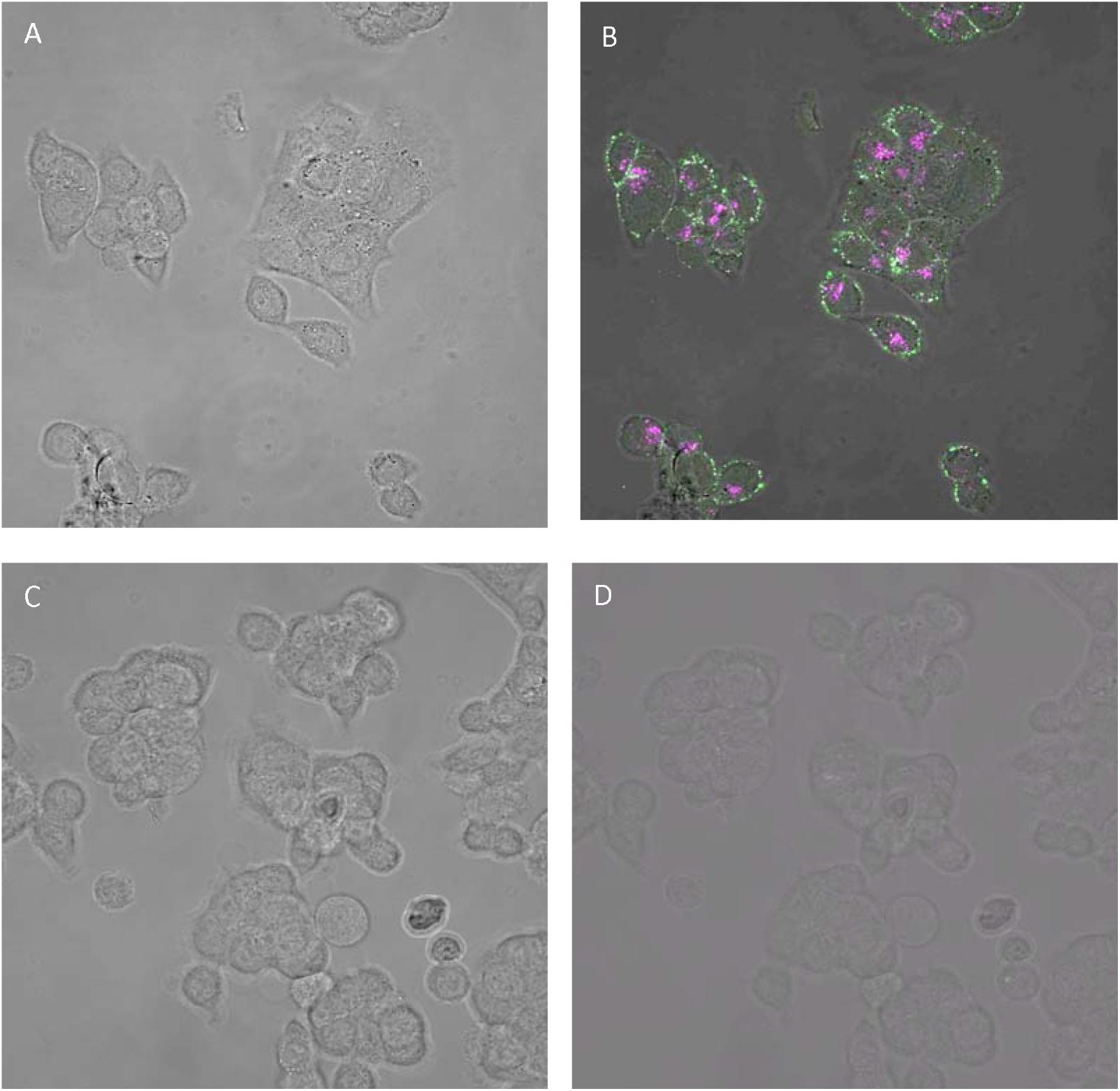
Confocal microscope images of the HCT116 cells. The cells are treated with loaded-nanoparticle in A and B, and empty nanoparticle in C and D. In A and C, images of the cells are shown. In B and D, images of the cells, alexa-fluo-647 signal (red) image and the bodipy-FL signal (green) image are merged together.

Thus, this experiment establishes that it is feasible to use nanoparticle-delivered fluorogenic nucleotide analog to monitor nucleotide-protein binding in live cells. Subsequently, we would be able to test whether the delivered BODIPY-FL-GTPγS can bind to the RAS proteins *in vivo* in HCT116 cells and activate its fluorescent activity, enabling a fluorescence-based RAS activity analysis method.

A good experimental method must be able to detect dynamic fluctuation of the targeted activity amid relevant cellular processes. Fortunately, one such process is available for testing whether our effort can achieve this capacity. It is well-known that serum starvation reduced RAS activity and that transient RAS activation would occur upon subsequent serum re-activation of the starved cells (34). The transient activation occurs quickly, within 20 minutes (34-36), providing a convenient testing opportunity for us. Thus, we monitored RAS-associated fluorescent activity during this process in nanoparticle-treated cells.

Our experimental workflow is shown in Figure 5. The approach is RAS immno-precipitation, followed by fluorescence analysis as described in Materials and Methods, with the results shown in figure 6. Briefly, HCT116 cells were plated into (1 × 10^6^ cells per dish), and grown in, 100 mm^2^ dishes. At log-phase, the cells were switched either to fresh regular medium or to medium without serum for serum starvation. Next day, the cells were incubated with BODIPY-GTP-γ-S-loaded LCP NPs for 4 h at an equivalent concentration of 10 μM. For serum-starved cells, 10% FBS was added to the dishes to re-activate the cells. At specified time points, the cells were harvested for cell lysate extraction. We pre-cleaned the cell lysates with a control antibody. The cell lysates were then aliquoted. One aliquot was subjected to immune-precipitation analysis with the pan RAS monoclonal antibody. We then quantified the fluorescent activities co-precipitated with the RAS proteins. Not surprisingly, the activity was lower in serum-starved cells; and a transient peak was observed at 15 minutes after serum re-activation (Fig. 6A). A Western blot analysis of another aliquot of the cell lysates showed that the RAS protein expression level remained constant throughout the process (Fig. 6B). Thus, the fluctuation of the fluorescent activity should be a reflection of the underlying regulation of RAS signaling activity during this process.

**Figure 5.**
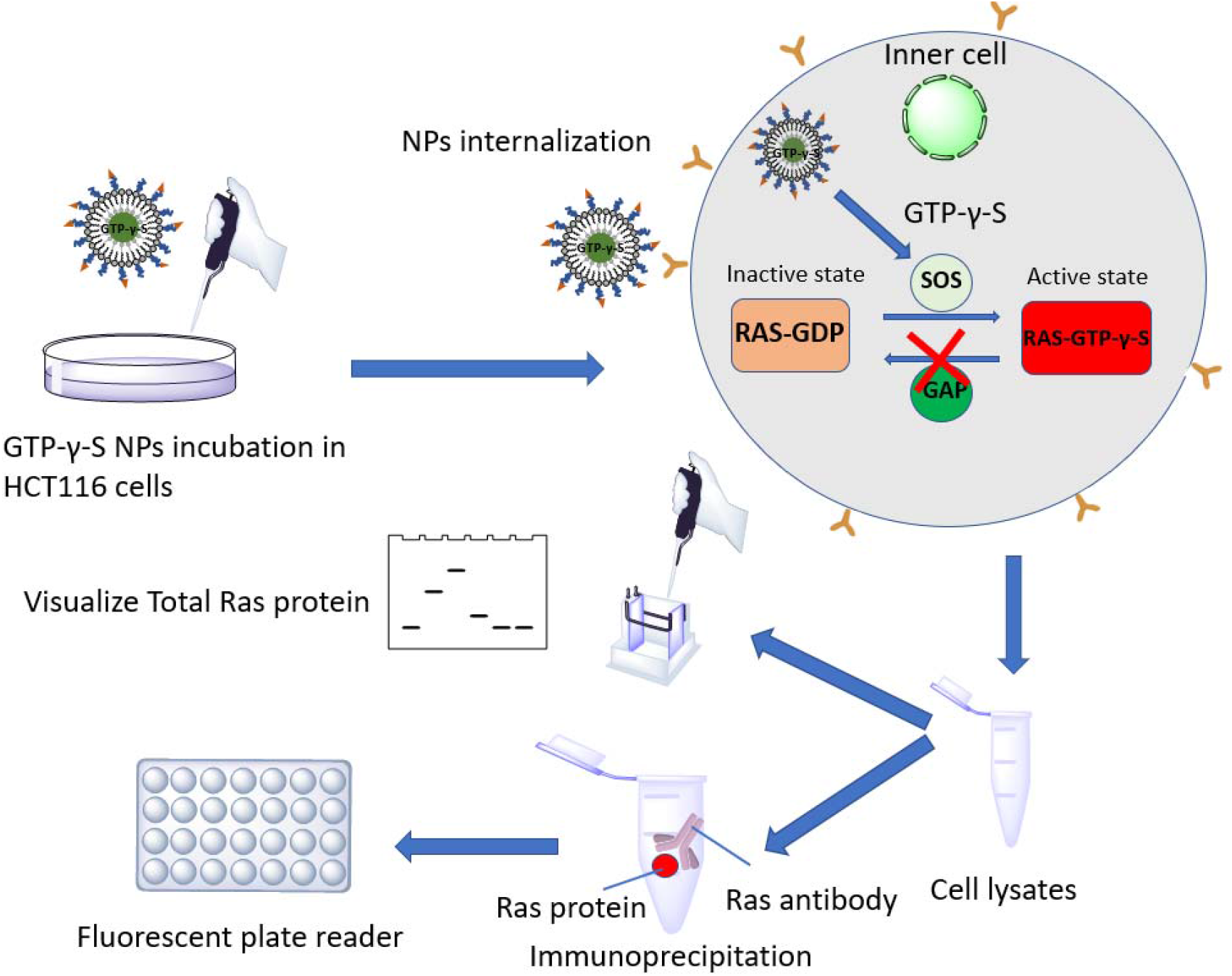
Our experimental strategy. Briefly, the cells were treated with the LCP nanoparticle. The cells were lysed, and the cell lysates were pre-cleaned with the control antibody and aliquoted. One aliquot was used for immune-precipitation and quantification of RAS-associated fluorescent activity, and another for Western blot analysis.

**Figure 6.**
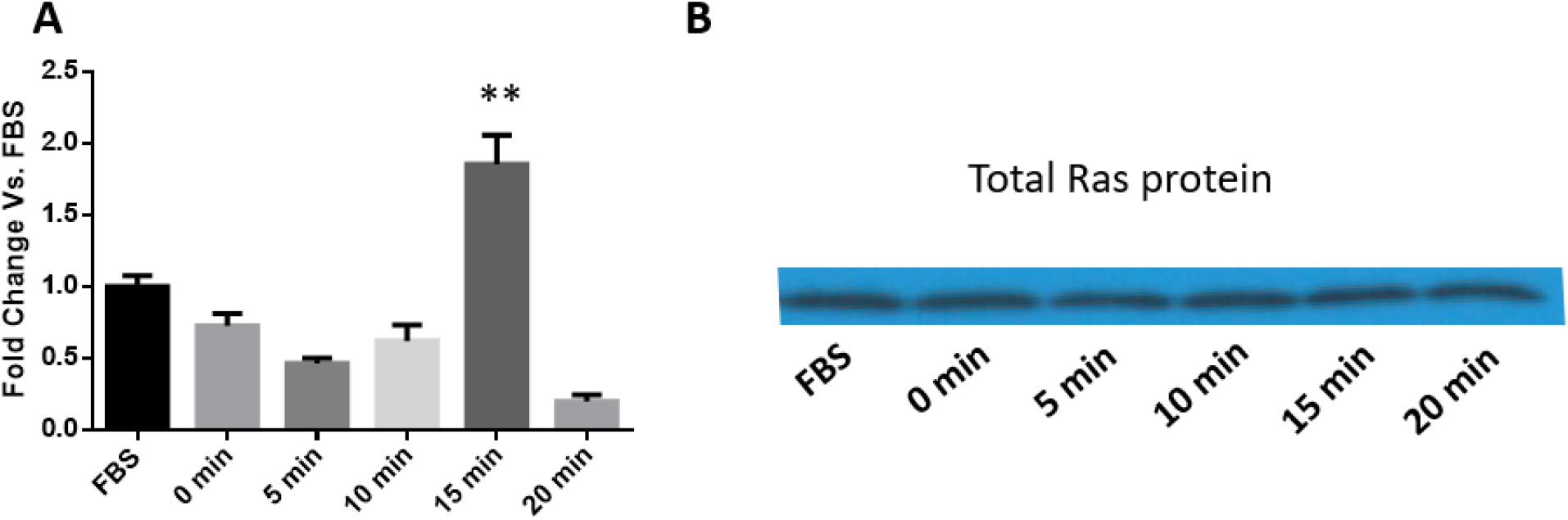
Analysis of cellular regulation of RAS via *in vivo* RAS-binding by delivered BODIPY-FL-GTPγS in HCT116 cells. (A) Dynamic fluctuation of RAS-associated fluorescent activity upon serum starvation, and at specified time points during subsequent serum reactivation, of the cells. The cells were lysed. Cell lysates were pre-cleaned with a control antibody and aliqoted. One aliquot set of the cleaned cell lysates were used for RAS immune-precipitation. The fluorescent activities co-precipitated with RAS were quantified and shown (**: *p* < 0. 01 in a t-test versus non-starved (FBS) cells). (B) Western blot analysis of another aliquot set of the pre-cleaned cell lysates with the RAS antibody, in order to illustrate equal RAS protein expression levels. The results shown in A and B are from one of 5 independent repeats of the whole experimental procedure; the same pattern was observed in all the repeats.

Taken together, the results suggested that the LCP nanoparticle was capable of intracellular delivery of sufficient amount of BODIPY-FL-GTPγS. Delivered BODIPY-FL-GTPγS molecules were able to compete with endogenous GTP for RAS binding, activating their fluorescent activity. The fluorescent activity, in turn, acted as a reporter of GTP binding to the RAS proteins. In a word, a RAS analysis method based on *in vivo* GTP binding activity was established.

## 4. Discussion

Many experimental protocols in biochemistry as well as molecular and cellular biology rely on biochemical analogs with easily quantifiable reporter activities as the analysis probes. Often, these protocols are rendered *in vitro* due to the lack of sufficient cell membrane permeability of these analysis probes. For examples, the analyses of RAS and other G-proteins that use GTP analogs as probes has to be executed *in vitro* with cell lysate, since GTP analogs are cell impermeable by themselves. In this study, we attempted to overcome this issue via nanoparticle-mediated intracellular delivery of a GTP analog with conditional fluorescent activity. The strategy should be applicable to many other experimental protocols.

Our approach currently requires an immune-precipitation step after *in vivo* binding of delivered BODIPY-FL-GTPγS to the RAS proteins. This is because of our inability to distinguish RAS GTP binding from other GTP binding events, which are numerous in the cells. For instance, GTP is also used as the energy source during the elongation stage of translation and, thus, binds to proteins of the translation machinery (37). As another example, there are many non-RAS G-proteins, which are often coupled to cell surface receptors (38). This necessitated lysing the cells, followed by RAS antibody immune-precipitation to isolate the RAS-specific GTP binding activity. Perhaps, the chemical genetic approach developed by Dr. Kevan Shokat for protein kinase substrate identification can be borrowed to address this issue (39). Briefly, in the Shokat approach, the gatekeeper amino acid residue in the ATP binding pocket of the protein kinase is mutated into a glycine to enlarge the pocket, so that the mutant protein kinase can accommodate a bulky ATP analog and use it in its kinase reaction. Consequently, the kinase reaction of the kinase of interest is distinguished from those of other protein kinases (more than 500) in the human kinome. A similar pair of RAS mutant and cognate GTP analogue, if successfully created, will be able to distinguish RAS GTP binding from other GTP binding events, enabling a fully *in vivo* RAS analysis method.

This strategy of using nanoparticle for intracellular probe delivery should be applicable to many other signal transduction analysis protocols. In addition to GTP, other nucleotides act frequently as allosteric regulators of key signaling proteins. For example, cyclic AMP (cAMP) is required for the activation of many proteins, such as protein kinase A (PKA) and the guanine nucleotide exchange factor (GEF) EPAC1 and EPAC2 (40); ADP and AMP activates the AMP-activated protein kinase (AMPK) (41). LCP nanoparticle should be capable of intracellular delivery of the analogs of these nucleotides and enable an *in vivo* version of the analysis protocols for the corresponding proteins.

Moreover, allosteric regulation by small chemicals occurs ubiquitously, *i*.*e*., in many other cellular functional domains, in the cells. First, the nucleotides are also frequent allosteric regulators of proteins with a wide range of biochemical functions. For instance, AMP activates phosphofructokinase 1 (PFK1), the enzyme at the rate-limiting step of the glycolysis pathway, and ATP competes with AMP to reverse the activation (42); and AMP activates glycogen phosphorylase b (PYGB) (43). Second, other types of chemicals also act as allosteric regulators. Among numerous such instances, Fructose 2,6-bisphosphate (Fru-2,6-P2) also regulates the PFK1 protein (44); the inhibition of 3-phosphoglycerate dehydrogenase (PHGDH) by serine; and the inhibition and activation of threonine deaminase by isoleucine and valine, respectively (45). In case the analogs of these allosteric regulators are used in the analysis of the cognate proteins, the experimental protocol will be a good candidate for *in vivo* improvement via nanoparticle-mediated intracellular delivery of the analogs. Likely, in order to achieve optimal delivery efficiency, different nanoparticles will be needed for different types of allosteric regulatory chemicals. And there are a wide variety of nanoparticles to choose from.

In summary, we tackled the issue of cell impermeability of key analysis probes, which rendered *in vivo* execution of the cognate experimental protocols impossible. In this initial effort, the LCP nanoparticle-mediated intracellular GTP analog delivery enabled an *in vivo* RAS analysis. Moreover, such allosteric regulation is ubiquitous in the cells. It is our firm belief that the principle and approach are applicable to many other protocols, opening up tremendous research opportunities.

## 5. Acknowledgments

This work was supported by National Institutes of Health (NIH) grants R15GM122006 to DW and U54CA198999 to LH.

## Notes

### Competing Interest Statement

The authors have declared no competing interest.

### Summary of Updates

Add one figure

## References

1. Core, L.J., Waterfall, J.J. and Lis, J.T. (2008) Nascent RNA sequencing reveals widespread pausing and divergent initiation at human promoters. Science, 322, 1845–1848.

2. Arkin, M.R., Connor, P.R., Emkey, R., Garbison, K.E., Heinz, B.A., Wiernicki, T.R., Johnston, P.A., Kandasamy, R.A., Rankl, N.B. and Sittampalam, S. (2004) In Sittampalam, G. S., Grossman, A., Brimacombe, K., Arkin, M., Auld, D., Austin, C. P., Baell, J., Bejcek, B., Caaveiro, J. M. M., Chung, T. D. Y.et al. (eds.), Assay Guidance Manual, Bethesda (MD).

3. Tour, O., Adams, S.R., Kerr, R.A., Meijer, R.M., Sejnowski, T.J., Tsien, R.W. and Tsien, R.Y. (2007) Calcium Green FlAsH as a genetically targeted small-molecule calcium indicator. Nat Chem Biol, 3, 423–431.

4. Cubitt, A.B., Heim, R., Adams, S.R., Boyd, A.E., Gross, L.A. and Tsien, R.Y. (1995) Understanding, improving and using green fluorescent proteins. Trends Biochem Sci, 20, 448–455.

5. Heim, R., Cubitt, A.B. and Tsien, R.Y. (1995) Improved green fluorescence. Nature, 373, 663–664.

6. Ni, Q. and Zhang, J. (2010) Dynamic visualization of cellular signaling. Adv Biochem Eng Biotechnol, 119, 79–97.

7. Algar, W.R., Hildebrandt, N., Vogel, S.S. and Medintz, I.L. (2019) FRET as a biomolecular research tool - understanding its potential while avoiding pitfalls. Nat Methods, 16, 815–829.

8. Herzog, V.A., Reichholf, B., Neumann, T., Rescheneder, P., Bhat, P., Burkard, T.R., Wlotzka, W., von Haeseler, A., Zuber, J. and Ameres, S.L. (2017) Thiol-linked alkylation of RNA to assess expression dynamics. Nat Methods, 14, 1198–1204.

9. Muhar, M., Ebert, A., Neumann, T., Umkehrer, C., Jude, J., Wieshofer, C., Rescheneder, P., Lipp, J.J., Herzog, V.A., Reichholf, B. et al. (2018) SLAM-seq defines direct gene-regulatory functions of the BRD4-MYC axis. Science, 360, 800–805.

10. Schofield, J.A., Duffy, E.E., Kiefer, L., Sullivan, M.C. and Simon, M.D. (2018) TimeLapse-seq: adding a temporal dimension to RNA sequencing through nucleoside recoding. Nat Methods, 15, 221–225.

11. Stephen, A.G., Esposito, D., Bagni, R.K. and McCormick, F. (2014) Dragging ras back in the ring. Cancer Cell, 25, 272–281.

12. Khanna, A., Lotfi, P., Chavan, A.J., Montaño, N.M., Bolourani, P., Weeks, G., Shen, Z., Briggs, S.P., Pots, H. and Van Haastert, P.J. (2016) The small GTPases Ras and Rap1 bind to and control TORC2 activity. Scientific reports, 6, 25823.

13. Do Heo, W. and Meyer, T. (2003) Switch-of-function mutants based on morphology classification of Ras superfamily small GTPases. Cell, 113, 315–328.

14. Takai, Y., Sasaki, T. and Matozaki, T. (2001) Small GTP-binding proteins. Physiological reviews, 81, 153–208.

15. Vetter, I.R. and Wittinghofer, A. (2001) The guanine nucleotide-binding switch in three dimensions. Science, 294, 1299–1304.

16. Korlach, J., Baird, D.W., Heikal, A.A., Gee, K.R., Hoffman, G.R. and Webb, W.W. (2004) Spontaneous nucleotide exchange in low molecular weight GTPases by fluorescently labeled gamma-phosphate-linked GTP analogs. Proc Natl Acad Sci U S A, 101, 2800–2805.

17. Schwartz, S.L., Tessema, M., Buranda, T., Pylypenko, O., Rak, A., Simons, P.C., Surviladze, Z., Sklar, L.A. and Wandinger-Ness, A. (2008) Flow cytometry for real-time measurement of guanine nucleotide binding and exchange by Ras-like GTPases. Anal Biochem, 381, 258–266.

18. Feig, L.A. and Cooper, G.M. (1988) Relationship among guanine nucleotide exchange, GTP hydrolysis, and transforming potential of mutated ras proteins. Molecular and cellular biology, 8, 2472–2478.

19. McEwen, D.P., Gee, K.R., Kang, H.C. and Neubig, R.R. (2001) Fluorescent BODIPY-GTP analogs: real-time measurement of nucleotide binding to G proteins. Analytical biochemistry, 291, 109–117.

20. Mittal, R., Ahmadian, M.R., Goody, R.S. and Wittinghofer, A. (1996) Formation of a transition-state analog of the Ras GTPase reaction by Ras· GDP, tetrafluoroaluminate, and GTPase-activating proteins. Science, 273, 115–117.

21. Willard, F.S., Kimple, A.J., Johnston, C.A. and Siderovski, D.P. (2005) A direct fluorescence-based assay for RGS domain GTPase accelerating activity. Analytical biochemistry, 340, 341–351.

22. Raepple, D., von Lintig, F., Zemojtel, T., Duchniewicz, M., Jung, A., Lubbert, M., Boss, G.R. and Scheele, J.S. (2009) Determination of Ras-GTP and Ras-GDP in patients with acute myelogenous leukemia (AML), myeloproliferative syndrome (MPS), juvenile myelomonocytic leukemia (JMML), acute lymphocytic leukemia (ALL), and malignant lymphoma: assessment of mutational and indirect activation. Ann Hematol, 88, 319–324.

23. Wilczewska, A.Z., Niemirowicz, K., Markiewicz, K.H. and Car, H. (2012) Nanoparticles as drug delivery systems. Pharmacol Rep, 64, 1020–1037.

24. Patra, J.K., Das, G., Fraceto, L.F., Campos, E.V.R., Rodriguez-Torres, M.D.P., Acosta-Torres, L.S., Diaz-Torres, L.A., Grillo, R., Swamy, M.K., Sharma, S. et al. (2018) Nano based drug delivery systems: recent developments and future prospects. J Nanobiotechnology, 16, 71.

25. Satterlee, A.B. and Huang, L. (2016) Current and Future Theranostic Applications of the Lipid-Calcium-Phosphate Nanoparticle Platform. Theranostics, 6, 918–929.

26. Zhang, Y., Kim, W.Y. and Huang, L. (2013) Systemic delivery of gemcitabine triphosphate via LCP nanoparticles for NSCLC and pancreatic cancer therapy. Biomaterials, 34, 3447–3458.

27. Zhang, Y., Peng, L., Mumper, R.J. and Huang, L. (2013) Combinational delivery of c-myc siRNA and nucleoside analogs in a single, synthetic nanocarrier for targeted cancer therapy. Biomaterials, 34, 8459–8468.

28. Dyawanapelly, S., Kumar, A. and Chourasia, M.K. (2017) Lessons Learned from Gemcitabine: Impact of Therapeutic Carrier Systems and Gemcitabine’s Drug Conjugates on Cancer Therapy. Crit Rev Ther Drug Carrier Syst, 34, 63–96.

29. Li, Y., Wu, Y., Huang, L., Miao, L., Zhou, J., Satterlee, A.B. and Yao, J. (2016) Sigma receptor-mediated targeted delivery of anti-angiogenic multifunctional nanodrugs for combination tumor therapy. Journal of Controlled Release, 228, 107–119.

30. Brattain, M.G., Marks, M.E., McCombs, J., Finely, W. and Brattain, D.E. (1983) Characterization of human colon carcinoma cell lines isolated from a single primary tumour. Br J Cancer, 47, 373–381.

31. Howell, G.M., Humphrey, L.E., Awwad, R.A., Wang, D., Koterba, A., Periyasamy, B., Yang, J., Li, W., Willson, J.K., Ziober, B.L. et al. (1998) Aberrant regulation of transforming growth factor-alpha during the establishment of growth arrest and quiescence of growth factor independent cells. J Biol Chem, 273, 9214–9223.

32. Jiang, W., Guo, Z., Lages, N., Zheng, W.J., Feliers, D., Zhang, F. and Wang, D. (2018) A Multi-Parameter Analysis of Cellular Coordination of Major Transcriptome Regulation Mechanisms. Sci Rep, 8, 5742.

33. Wang, D., Wang, T., Gill, A., Hilliard, T., Chen, F., Karamyshev, A.L. and Zhang, F. (2020) Uncovering the cellular capacity for intensive and specific feedback self-control of the argonautes and MicroRNA targeting activity. Nucleic Acids Res, 48, 4681–4697.

34. Giehl, K., Skripczynski, B., Mansard, A., Menke, A. and Gierschik, P. (2000) Growth factor-dependent activation of the Ras-Raf-MEK-MAPK pathway in the human pancreatic carcinoma cell line PANC-1 carrying activated K-ras: implications for cell proliferation and cell migration. Oncogene, 19, 2930–2942.

35. Kranenburg, O., Verlaan, I. and Moolenaar, W.H. (2001) Regulating c-Ras function. cholesterol depletion affects caveolin association, GTP loading, and signaling. Curr Biol, 11, 1880–1884.

36. Augsten, M., Pusch, R., Biskup, C., Rennert, K., Wittig, U., Beyer, K., Blume, A., Wetzker, R., Friedrich, K. and Rubio, I. (2006) Live-cell imaging of endogenous Ras-GTP illustrates predominant Ras activation at the plasma membrane. EMBO Rep, 7, 46–51.

37. Wallin, G., Kamerlin, S.C. and Aqvist, J. (2013) Energetics of activation of GTP hydrolysis on the ribosome. Nat Commun, 4, 1733.

38. McCudden, C.R., Hains, M.D., Kimple, R.J., Siderovski, D.P. and Willard, F.S. (2005) G-protein signaling: back to the future. Cell Mol Life Sci, 62, 551–577.

39. Hertz, N.T., Wang, B.T., Allen, J.J., Zhang, C., Dar, A.C., Burlingame, A.L. and Shokat, K.M. (2010) Chemical genetic approach for kinase-substrate mapping by covalent capture of thiophosphopeptides and analysis by mass spectrometry. Curr Protoc Chem Biol, 2, 15–36.

40. Serezani, C.H., Ballinger, M.N., Aronoff, D.M. and Peters-Golden, M. (2008) Cyclic AMP: master regulator of innate immune cell function. Am J Respir Cell Mol Biol, 39, 127–132.

41. Mihaylova, M.M. and Shaw, R.J. (2011) The AMPK signalling pathway coordinates cell growth, autophagy and metabolism. Nat Cell Biol, 13, 1016–1023.

42. Webb, B.A., Forouhar, F., Szu, F.E., Seetharaman, J., Tong, L. and Barber, D.L. (2015) Structures of human phosphofructokinase-1 and atomic basis of cancer-associated mutations. Nature, 523, 111–114.

43. Walcott, S. and Lehman, S.L. (2007) Enzyme kinetics of muscle glycogen phosphorylase b. Biochemistry, 46, 11957–11968.

44. Hue, L. and Rider, M.H. (1987) Role of fructose 2,6-bisphosphate in the control of glycolysis in mammalian tissues. Biochem J, 245, 313–324.

45. Grant, G.A. (2018) D-3-Phosphoglycerate Dehydrogenase. Front Mol Biosci, 5, 110.

